# LORA, Lipid Over-Representation Analysis based on structural information

**DOI:** 10.1101/2023.06.02.543363

**Authors:** Michaela Vondrackova, Dominik Kopczynski, Nils Hoffmann, Ondrej Kuda

## Abstract

With the increasing number of lipidomic studies, there is a need for efficient and automated analysis of lipidomic data. One of the challenges faced by most existing approaches to lipidomic data analysis is lipid nomenclature. The systematic nomenclature of lipids contains all available information about the molecule, including its hierarchical representation, which can be used for statistical evaluation. The Lipid Over-Representation Analysis (LORA) web application (https://lora.metabolomics.fgu.cas.cz) analyzes this information using the Java-based Goslin framework, which translates lipid names into a standardized nomenclature. Goslin provides the level of lipid hierarchy, including information on headgroups, acyl chains, and their modifications, up to the ‘complete structure’ level. LORA allows the user to upload the experimental query and universe datasets, select a grammar for lipid name normalization, and then process the data. The user can then interactively explore the results and perform lipid overrepresentation analysis based on selected criteria. The results are graphically visualized according to the lipidome hierarchy. The lipids present in the most over-represented terms (lipids with the highest number of enriched shared structural features) are defined as Very Important Lipids (VILs). For example, the main result of a demo dataset is the information that the query is significantly enriched with ‘glycerophospholipids’ containing ‘acyl 20:4’ at ‘*sn*-2 position’. These terms define a set of VILs (e.g., PC 18:2/20:4;O and PE 16:0/20:4(5,8,10,14);OH). All results, graphs, and visualizations are summarized in a report. LORA is a tool focused on the smart mining of epilipidomics datasets to facilitate their interpretation at the molecular level.

## INTRODUCTION

Recent advances in analytical techniques and their routine use in screening pipelines push forward our ability to provide a full structural characterization of lipid species in complex biological matrices. Stereospecifically numbered (*sn*) position of acyl/alkyl chain on the glycerol backbone, double bond position and stereochemistry within acyl/alkyl chains, and configuration of chiral centers can be assigned using a combination of advanced separation and ion activation techniques.^1^ The emerging challenge is to correctly and systematically annotate the lipid species ^2^ and consider the high structural diversity of modified lipid species, generally referred to as the ‘epilipidome’.^3^ Correctly annotated and standardized lipid datasets represent an information source for further FAIR data mining.

Over-Representation Analysis (ORA) is a simple statistical method that determines whether an a priori-defined set of variables is more present (over-represented) in a subset of variables than would be expected by chance. Two main bioinformatics tools for over-representation analysis of lipidomics datasets are available: 1) LION^4^, a lipid ontology tool that associates >50,000 lipid species to biophysical, chemical, and cell biological features; and 2) Lipid Mini-On^5^, an open-source tool that performs lipid enrichment analyses and visualizations of lipidomics data. Both tools use custom-defined lipid databases, specific nomenclatures, and parsing functions to mine data from lipid names. However, these tools lack the power to exploit the hierarchical nature of lipid structures and leave the data unmined.

The goal of the LORA project was to build a bioinformatics tool based on Goslin, a systematic grammar-based lipid library, to facilitate the statistical evaluation of lipid structural information and to support international standardization of lipidomics nomenclature.^2,6,7^

## METHODS

### Lipid Identifiers and Goslin

Bioinformatic tools for lipid ORA require lipid names or database identifiers. We used jGoslin, the Java implementation of Goslin^7^, which parses the submitted lipid names and translates them into a normalized hierarchical representation (Table S1). jGoslin supports lipid names based on LIPID MAPS, SwissLipids, HMDB, and Shorthand nomenclature. Lipid classes implemented in Goslin are periodically updated as new lipid classes are discovered.^7^ Epilipidome nomenclature can be converted to a compatible format by LipidLynxX.^8^

### Datasets

The lipid datasets used in this work refer to four published datasets from lipidomic studies in humans: 1) DEMO 1: Cachexia, body weight stable vs. cachectic patients (human epicardial adipose tissue)^9^; 2) DEMO 2: AdipoAtlas, obese vs. lean patients (human white adipose tissue)^10^; 3) DEMO 3: Lipid Mini-On demo dataset (human lung endothelial lipidome)^5^; 4) technical DEMO 4: oxidized membrane lipids (human platelets) at ‘Complete structure level’, combined with phospholipids at ‘Structure defined’ level (mouse liver), and Goslin performance test file.^6,11,12^ A list of query lipids, the whole lipidome (universe), LORA manual, and the LORA report for each dataset is attached as Supplemental Material and available in the LORA web application.

### Implementation of ORA

Two enrichment tests were used to perform ORA: 1) Fisher exact test (scipy.stats.fisher_exact) with ‘two-sided’, ‘less’, ‘greater’ alternatives; and 2) Hypergeometric test (scipy.stats.hypergeom) from SciPy^13^. Multiple comparisons were adjusted using Bonferroni, Holm-Bonferroni, or Benjamini-Hochberg procedure (the classic False Discovery Rate, FDR).^14,15^ The ORA was performed at each level of the nomenclature hierarchy and on structural features provided by Goslin to correct for the set size effect.^16^ Additional parameters (grouped number of carbon atoms per acyl chain: less than 16, 16–18, and more than 18; and the number of double bonds: 0, saturated; 1, monounsaturated; 2 and more, polyunsaturated) were included to facilitate biological interpretation of the data. UpSet plots provide visualization of term intersections.^17^ The visualization of nomenclature levels was implemented via Biopython.^18^ The lipidome was converted to phyloXML format, an XML language designed to describe phylogenetic trees (or networks) and associated data^19^, and the pseudo-phylogenetic distances (radii of the spheres/levels) were optimized for the hierarchical structure of the lipidome. Only the CATEGORY and CLASS levels are labeled to optimize space use and prevent overlapping labels. The Cytoscape network was built to visualize the quantitative data and statistics.^20^

## RESULTS

### LORA, a web-based tool for Lipid Over-Representation Analysis

We developed LORA as a web-based interactive tool using Python 3.10, Dash, and Plotly packages.^21,22^ The application uses a tabular layout. The query and universe lists of lipids are uploaded, processed, and parsed via Goslin. Users can change the ORA parameters, perform the analysis, interact with the plots, and download the report (Figure 1). LORA is available as a web service (https://lora.metabolomics.fgu.cas.cz), and the source code is hosted at GitHub https://github.com/IPHYSBioinformatics/LORA under the terms of liberal open source licenses.

**Figure 1.**
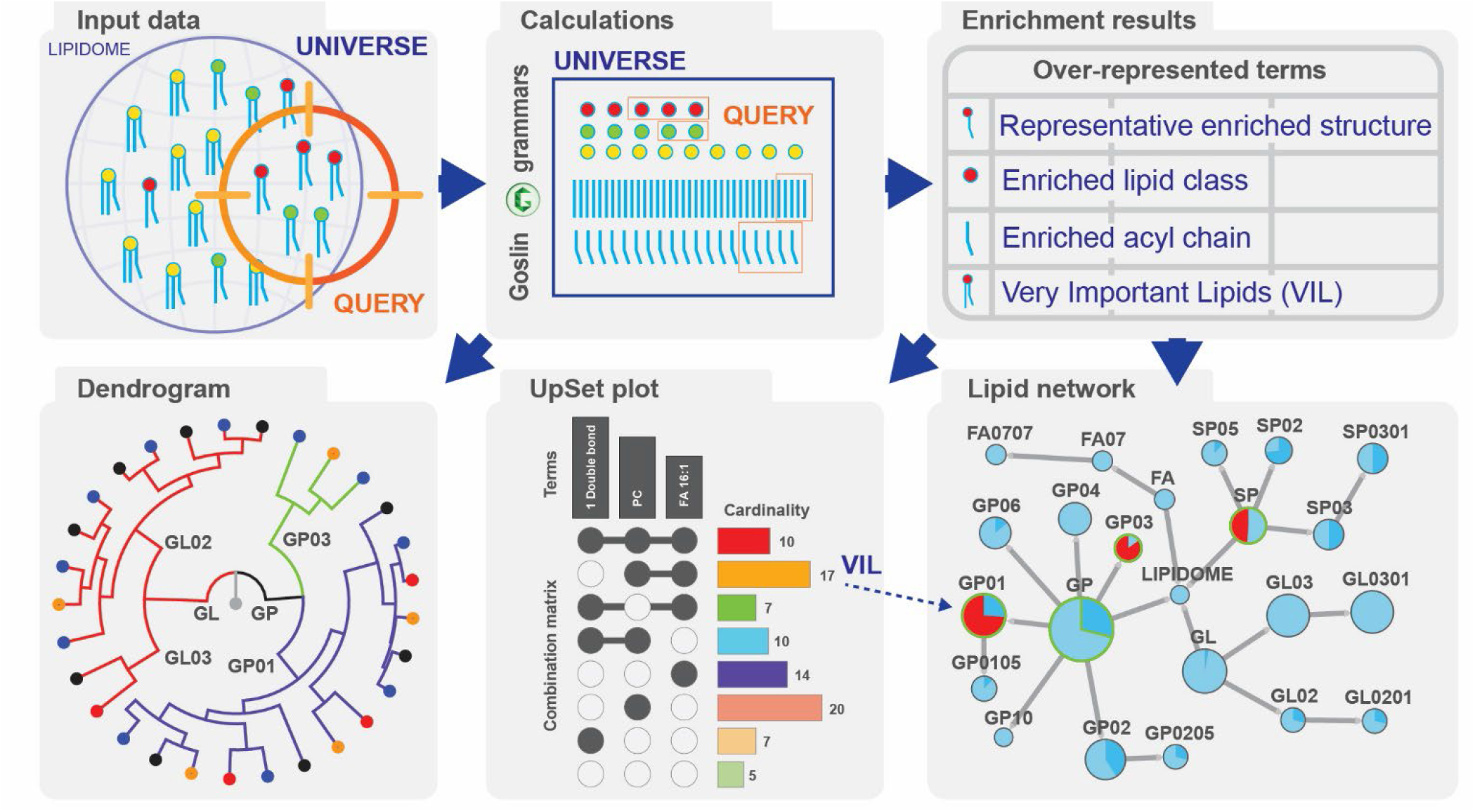
LORA pipeline. User data (universe and query lipid names) are processed using Goslin, the lipidome is visualized as a dendrogram, and LORA is performed. Enrichment results are summarized in a table, which is further processed to a lipid network and UpSet plot.

The primary output of ORA is a set of over-represented terms. The terms are defined based on the nomenclature levels and structural characteristics of the lipidome parsed by Goslin, e.g., ‘Total number of carbon atoms’, ‘Fatty Acid #1 *sn* position’, Total #O, etc.

LORA is the first tool that uses smart text mining and extracts all available structural information from the provided lipid identifier. Therefore, enrichment based on double bond positions (location and conformation), bond type (ester, ether), and modifications (oxidized, cyclized) can be calculated. Human adipose tissue lipidomes (DEMO 1 and DEMO 2) demonstrate how LORA mines the information from widely used LC–MS lipidomic pipelines.^9,10^ Human lung lipidome dataset in DEMO 3 comes from Lipid Mini-On test files. Technical DEMO 4 dataset contains a collection of (modified) lipids defined up to ‘Complete structure’ to illustrate the potential use.

### Shared structural characteristics of over-represented lipids

The idea behind ORA is that we can infer a smaller set of structural characteristics for a set of significantly altered lipids, which reduce the dimensionality and define the essential features of query data. However, we can also take the sets of structural characteristics, explore their intersections, and find the lipid molecule(s) that best represent the set of significantly altered lipids. The lipids present in the most over-represented terms (lipids with the highest number of term intersections) are defined as Very Important Lipids (VILs). Table 1 and Figure 2 show the analysis of DEMO 4. Table 1 defines the seven overrepresented terms, and the UpSet plot in Figure 2 highlights the common patterns. The green cluster defines 13 lipids that share structural features in four over-represented terms (Figures 2B and 2C). For example, one output is: “The query lipids are statistically significantly enriched in glycerophospholipids containing 20:4 acyl at the *sn*-2 position”.

**Table 1.** Summary of over-represented terms from DEMO 4 calculated at alpha level 0.001 with FDR correction.

| Term (Group / Classifier) | Goslin Level | N<sub>Q</sub> Query | N<sub>U</sub> Universe | *p*-value | Odds Ratio | FDR |
| --- | --- | --- | --- | --- | --- | --- |
| Acyls 16:1 | MOLECULAR SPECIES | 13/68 | 18/556 | 4.30e-06 | 7.0646 | 0 |
| Acyls 20:4 | MOLECULAR SPECIES | 15/68 | 26/556 | 5.44e-06 | 5.7692 | 0 |
| Acyls 20:4 | SN POSITION | 13/68 | 19/556 | 6.63e-06 | 6.6804 | 0.0001 |
| Acyls 16:1 within Glycerophospholipids [GP] | MOLECULAR SPECIES | 13/68 | 17/448 | 2.36e-05 | 5.9925 | 0.0002 |
| Acyls 20:4 within Glycerophospholipids [GP] | MOLECULAR SPECIES | 15/68 | 24/448 | 2.86e-05 | 5.0000 | 0.0002 |
| Acyls 20:4 within Glycerophospholipids [GP] | SN POSITION | 13/68 | 18/448 | 3.64e-05 | 5.6465 | 0.0004 |
| Acyls 16:1 within Glycerophosphocholines [GP01] | MOLECULAR SPECIES | 11/26 | 14/148 | 1.00e-4 | 7.0190 | 0.0005 |

**Figure 2.**
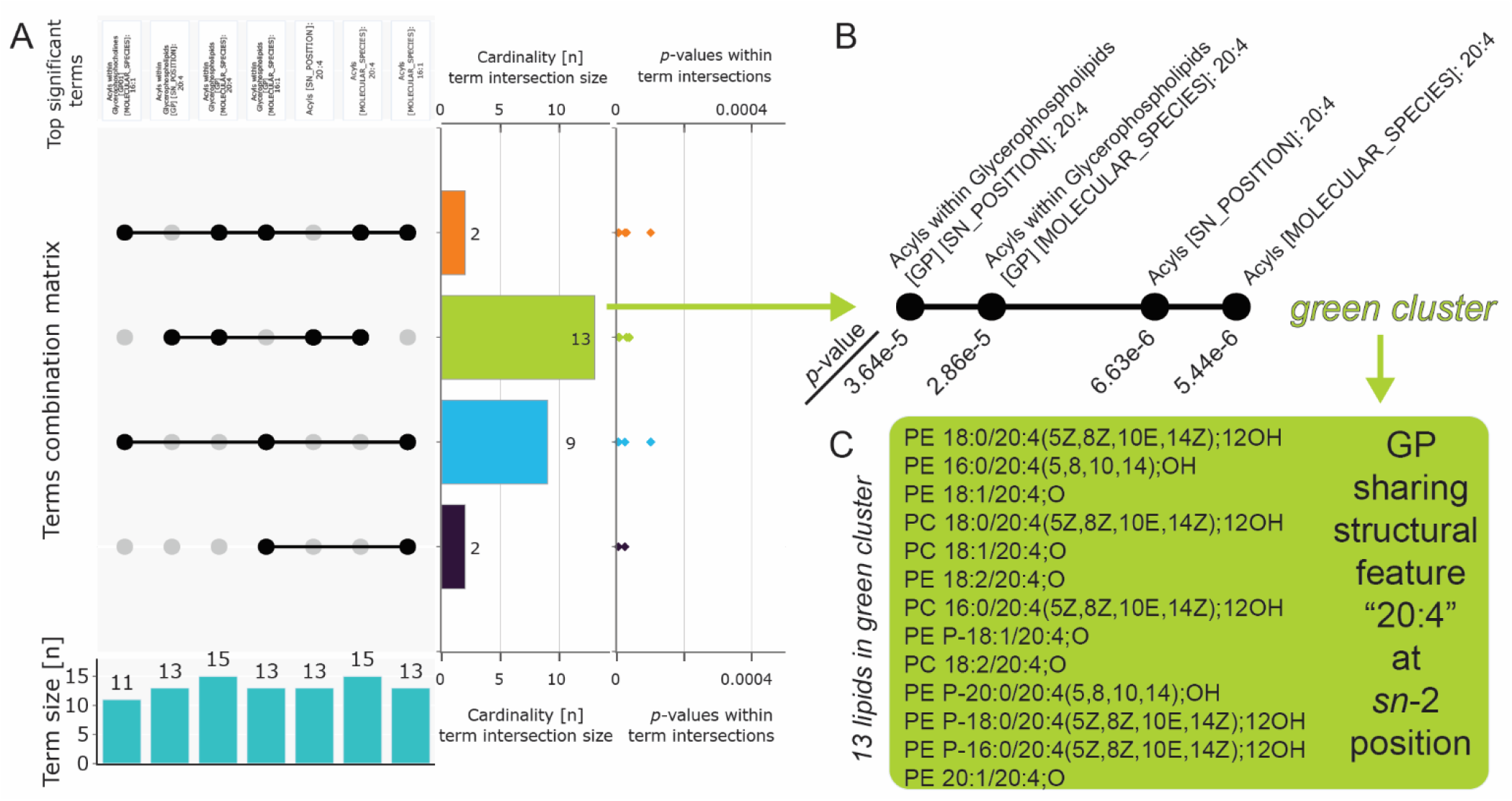
Interpretation of LORA results. A: UpSet plot showing the intersection of over-represented terms (structural characteristics) from Table 1. Cardinality is sorted according to the total number of term intersections. B: Intersection of 4 terms defining the green cluster. C: Green cluster containing 13 lipids which share a structural feature “20:4” at *sn*-2 position.

### UpSet plot visualization of term intersections

The major challenge in evaluating the over-represented structural characteristics is the enormous number of set intersections if the number of sets exceeds a reasonable threshold. In the case of one to three sets, the Venn or Euler diagrams, which visualize all possible logical relations between the sets, can be used. In the case of two to twenty sets, the UpSet plot provides a comprehensive visualization.^17^ The Upset plot helps identify enriched lipids’ main structural features and highlights the VILs in a graphical representation (Figure 2A). Connected black dots represent the intersection of the terms labeled on top. The term intersection size (cardinality bar plot) represents the number of lipids that have this specific set of structural features in common. The *p*-values belong to the particular lipids within individual terms (*n* lipids × *m* terms). The bar graph at the bottom shows how many lipids belong to the term. The plot is interactive, and a table showing all lipids within the specific intersection (Figure 2C) is generated upon clicking on the cardinality bar.

### Hierarchical tree visualization of the lipidome

Goslin provides information on the hierarchical level of the lipids in the dataset. We can visualize the information as a circular tree map (dendrogram) showing part-to-whole relationships and the level of structural details provided by the analytical method. The graph itself is interactive, and each lipid level can be explored via a tooltip (Figure S1A). The lipidome enrichment down to the LIPID MAPS SUBCLASS level can be visualized as an interactive network in Cytoscape, which allows the mapping of statistical data onto nodes and edges (Figure S1B).

### Report

LORA provides output in a zip archive containing a PDF report, parameter settings, over-representation analysis results, VIL table, UpSet plot, intersection tables, circular tree map, and the lipidome network in SVG, JPG, phyloXML, interactive HTML, XLSX, and Cytoscape formats, respectively.

### Limitations

Lipid modifications, including oxidation, nitration, or halogenation, represent a new level of nomenclature complexity that extremely expands the search space.^3^ Testing all structural characteristics up to ‘Complete structure level’ would require computational resources that would not balance information gain. Therefore, the application does not implement ‘rare’ features like the position of lipid modification or enantiomers. Lipid names in conflict with the hierarchical structure of shorthand notation (e.g., PC 16:1(7)_16:1(9)) will be converted to the closest valid level (e.g., PC 16:1_16:1). Goslin grammars (version 2) are currently limited to the most common set of lipid classes and do not consider multiple class categorization of lipids in LIPID MAPS (e.g., [FA01] Fatty Acid Conjugates and [FA02] Octadecanoids).^7^

## DISCUSSION AND CONCLUSION

Lipidomic analyses are usually performed to gain insight into system lipid metabolism, and ORA helps reduce the observation’s dimensionality. In contrast to transcriptomics, where the ORA results can be directly used in pathways analysis^16^, functional pathway schemes for lipids are largely unavailable. The major problems are 1) the inability to assign a lipid database identifier to experimentally generated information describing the lipid molecule at the level of enzyme-substrate specificity^2^; 2) the continuous remodeling of the head groups and acyl chains by many enzymes simultaneously; and 3) the lack of curated lipid pathways at various nomenclature levels (with a few exceptions^23,24^). Generalized databases like KEGG do not consider the structural diversity of lipids and often blur together multiple biological processes, thus compromising the biological interpretation. To overcome the problems, LORA builds on the Goslin standardization approach and known lipid structural characteristics provided by novel analytical techniques. Instead of relying on pre-defined schemes, LORA creates a set of overrepresented terms based solely on provided structural information and statistical tests. The user can either directly interpret this set or re-shape it by the UpSet plot to highlight the most common structural features and their representatives (lipid species).

The most common set visualization approach – Venn diagrams – does not scale beyond three or four sets. The UpSet plot, in contrast, is well suited for the quantitative analysis of data with more than three sets. When more than seven sets intersect, the advanced UpSet plots allowing aggregation & grouping should be used to reduce the dimensionality.^17^ We optimized the UpSet plot implemented in LORA for common lipidomics datasets. We limited the visualization to at most 13 terms because the calculation costs of all possible term intersections grow exponentially. Of note, the choice of lipid structural features, similar to the pathway database, used in ORA can have a much stronger effect on the enrichment results than the statistical corrections used in these analyses.^16^

LORA is a tool focused on the expanding technologies in (epi)lipidomics that allow more precise identification of lipid structures. Routine use of supercritical fluid chromatography, ion mobility spectrometry, ion-molecule reactions, or derivatization techniques to specifically target double bond positions will provide further levels of detail to lead towards the full structural characterization of lipids. LORA mines this information-rich dataset and helps interpret lipid structural features and over-represented terms using visualization tools. It is the next step toward understanding lipidomic datasets at the molecular level.

## ASSOCIATED CONTENT

### Supporting Information

The Supporting Information is available free of charge on the ACS Publications website.

- Manual (PDF)
- Supplementary figures (PDF)
- Reports generated from DEMO datasets (ZIP)
- Source code (ZIP)

## Author Contributions

Conceptualization: O.K.; Data curation: M.V. and O.K.; Formal analysis: M.V., D.K. and N.H.; Funding acquisition: O.K.; Investigation: M.V. and O.K.; Methodology: M.V. and O.K.; Project administration: O.K.; Resources: D.K., N.H. and O.K.; Software: M.V., D.K., N.H. and O.K.; Supervision: O.K.; Validation: D.K. and N.H.; Visualization: M.V. and O.K.; Writing – original draft: M.V. and O.K.; Writing - review & editing: M.V., N.H. and O.K.

### Notes

The authors declare no competing financial interests.

## ACKNOWLEDGMENT

Supported by the project National Institute for Research of Metabolic and Cardiovascular Diseases (Programme EXCELES, ID Project No. LX22NPO5104) – Funded by the European Union – Next Generation EU, Ministry of Health [NV19-02-00118], the Czech Academy of Sciences [Lumina Quaeruntur LQ200111901] and COST Action CA19105 - Pan-European Network in Lipidomics and EpiLipidomics (EpiLipidNET), supported by COST (European Cooperation in Science and Technology).

---

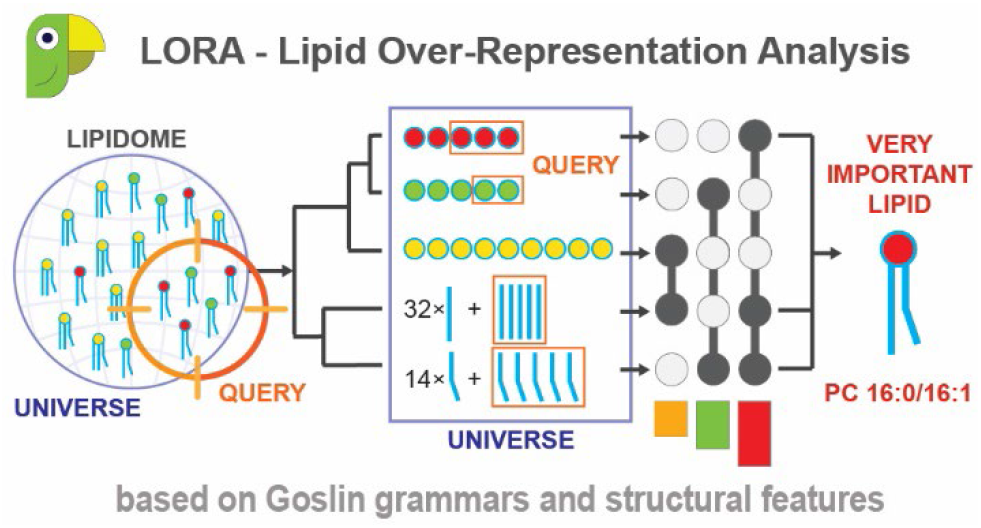

