## Supplemental data for "LORA, Lipid Over-Representation Analysis based on structural information"

#### Lipidome visualization

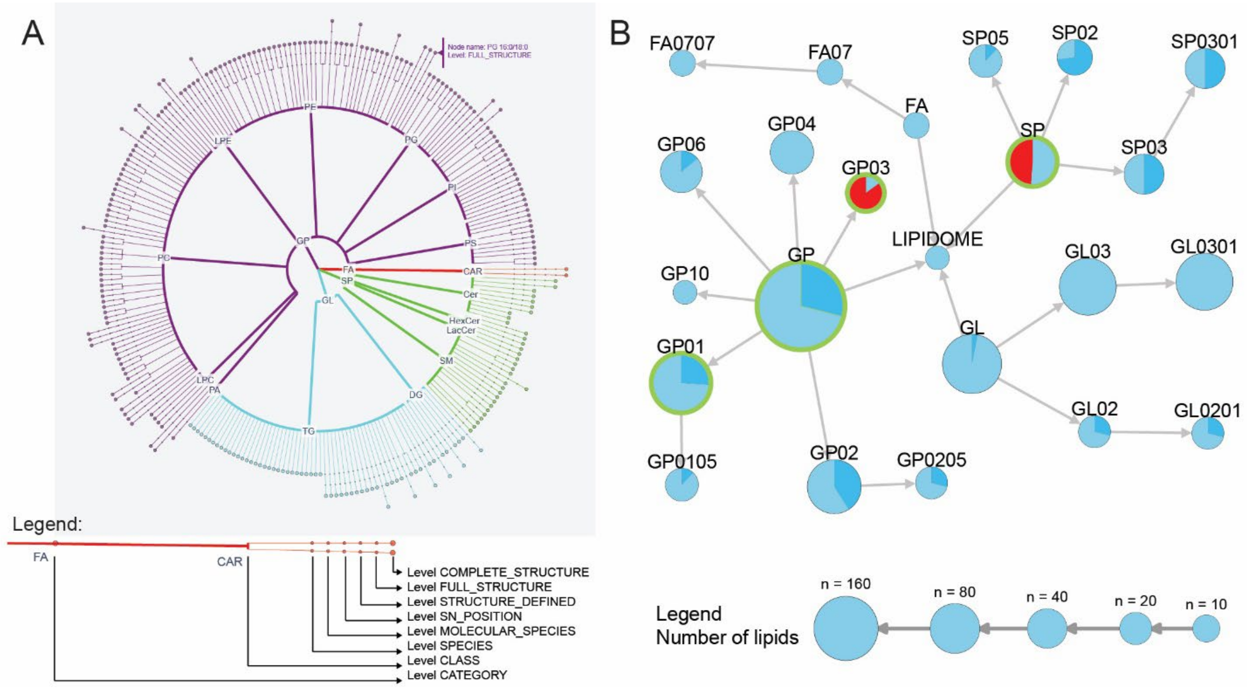

Figure S1: **Lipidome visualization**. A: Hierarchical tree based on Goslin levels for each lipid molecule. B: Lipid network highlighting the number of species and related LORA statistics. Abbreviations of lipid categories, classes and subclasses follow LIPIDMAPS nomenclature and SHORTHAND notation [1].

#### Comparison of LORA with other software tools

Compared to Lipid Mini-On [2], LORA provides more rigorous results by systematically parsing structural information and applying standardized lipid nomenclature rules (see Figure S1A and Table S1). While Lipid Mini-On generates an enrichment network of lipids and terms and stacked bar plots for the classifiers, LORA generates an UpSet plot, a hierarchical lipidome tree, and a structured lipid network with statistically significant values. Therefore, the tools are complementary when provided with a valid list of lipids. In addition, several features improve the performance of LORA over Lipid Mini-On: 1) LORA accepts multiple lipid nomenclature dialects thanks to jGoslin conversion; 2) new LM categories and updates are propagated to LORA via Goslin, and no manual database update is required; and 3) ORA is performed at a specific nomenclature level using all available structural information.

Compared to LION [3], LORA offers the same advantages as Lipid Mini-On, whereas LION contains a hard-coded database of generalized biophysical, chemical, and cell biological properties (MeSH terms) associated with lipids. The main drawback of LION is its outdated database (v. 2020.07.14), which was updated prior to the publication of the Lipid Shorthand Notation.

Alternatively, bioinformatics tools for ORA of general metabolomics data, such as MetaboAnalyst [4], can be used. However, such tools ignore the lipid structural information and work only with general identifiers.

Epilipidomics datasets containing detailed information about lipid modifications can be pre-processed by LipidLinxX [5] and converted into a nomenclature format compatible with currently supported Goslin grammars.

### Goslin lipid structural hierarchy

**Table S1**. Structural hierarchy representation of PE 16:1(6Z)/16:0;5OH[R],8OH;3oxo). LM: LIPID MAPS, HG: Head Group, FA: Fatty Acyl. Adapted from [6]

| Level | Name | Description |
|---|---|---|
| Category (LM) | Glycerophospholipids (GP) | Lipid category |
| Class (LM) | Glycerophosphoethanolamine (PE) GP02 | Lipid class |
| Species (LM Subclass) | Phosphatidylethanolamine, PE 32:2;O3 | HG, FA summary, two double bond equivalents, three oxidations |
| Molecular species | PE 16:1_16:1;O3 | HG, two FAs, two double bond equivalents, three oxidations |
| *sn*-Position | PE 16:1/16:1;O3 | HG, SN positions, here: for FA1 at *sn*-1 and FA2 at *sn*-2, two double bond equivalents, three oxidations |
| Structure defined | PE 16:1(6)/16:1;(OH)2;oxo | HG, SN positions, here: for FA1 at *sn*-1 and FA2 at *sn*-2, three oxidations and unspecified stereo configuration (6) on FA1 |
| Full structure | PE 16:1(6Z)/16:1;5OH,8OH;3oxo | HG, SN positions, here: for FA1 at *sn*-1 and FA2 at *sn*-2, positions for oxidations and stereo configuration (6Z) on FA1 |
| Complete structure | PE 16:1(6Z)/16:0;5OH[R],8OH;3oxo | HG, SN positions, here: for FA1 at *sn*-1 and FA2 at *sn*-2, positions for oxidations and stereo configuration ([R]) and double bond position and stereo configuration (6Z) on FA1 |
